# Dynamic cancer dormancy and awakening emerge from tumor microenvironment feedback in a minimal theoretical model

**DOI:** 10.64898/2026.04.14.718509

**Authors:** Guillermo Yáñez Feliú, Giacomo Rossato, Angelo Valleriani, Amaia Cipitria

## Abstract

Cancer cell dormancy is characterized by late relapse and therapy resistance, yet the mechanisms that awaken dormant cells remain poorly understood. The tumor microenvironment has emerged as a key driver of these state transitions. Here we present a theoretical framework based on evolutionary game theory in which interactions between cancer and host cells are coupled explicitly to a changing tumor microenvironment. Cancer cells produce a conditioning factor that is cleared by the microenvironment and tolerated only up to a threshold. Through this conditioning factor, the microenvironment feeds back on cancer–host interactions and reshapes their competitive balance. Unlike a model with fixed interactions, this feedback allows dormancy and awakening to emerge as dynamic outcomes of microenvironmental change. We show that this minimal coupling is sufficient to generate distinct long-term regimes. Across these regimes, feedback generates thresholds and history dependence, so that the same cancer population can follow different fates depending on whether the microenvironment is already primed. Our framework reduces these dynamics to experimentally and biologically interpretable parameters linked to conditioning factor production, clearance, tolerance, and microenvironment-dependent changes in cancer–host competition. More broadly, it provides a quantitative basis for testing how collective microenvironmental feedback shapes cancer dormancy and awakening, and for designing experiments to uncover the mechanisms that awaken dormant cancer cells.

## 1 Introduction

Metastatic progression accounts for most cancer mortality [1–3]. Metastatic relapse can occur long after apparent clinical resolution [4, 5]. Although cancer dormancy has been described across multiple tumor types, breast cancer provides a particularly important and well-studied context because it remains the leading cause of cancer death in women worldwide [6, 7].This recurrence is often linked to disseminated cancer cells (DCCs) that persist in distant microenvironments for extended periods in a non-proliferative or weakly proliferative state [8,9]. Such persistence can be history-dependent, as early disseminated cells may prime the microenvironment through secreted signals, modify it in ways that make later arrivals more likely to survive and progress [9,10]. Breast cancer DCCs show organ-specific colonization patterns, with metastases most frequently detected in bone, lung, liver, and brain [11, 12]. This long-lived DCC state, broadly referred to as cancer dormancy, underlies therapy resistance and late relapse, yet the mechanisms that trigger awakening from dormancy and subsequent metastatic outgrowth remain poorly understood [13–16].

The transition from dormancy to metastatic outgrowth is not immediate but is a multi-step process. Rather than treating dormancy as a static arrested state, an emerging view is that dormancy is a dynamic process that evolves across biological scales in space and time [17]. The clinically asymptomatic phase can involve single dormant DCCs, tumor mass dormancy where proliferation is balanced by cell death, and small active micrometastases that remain subclinical [10,15,17,18]. Transitions between these states are driven by both cell-intrinsic programs and fluctuating microenvironmental cues. A growing body of evidence indicates that the microenvironment strongly shapes dormancy, awakening, and metastatic outgrowth, and actively regulates transitions between cancer cell states [19–27].

Microenvironmental cues include biochemical and biophysical factors, including nutrient and oxygen availability [28], immune surveillance [29, 30], and extracellular matrix (ECM) properties [31, 32], among others. These cues can influence survival and quiescence programs, constrain proliferation, and modulate the ability of DCCs to colonize and expand [33]. In particular, ECM composition, organization, and mechanics can actively regulate dormancy and relapse [34]. For example, cell–matrix interactions have been shown to regulate dormancy in cancer stem-like populations [35]. Recent *in vivo* studies show that dormancy and outgrowth are regulated by niche-specific ECM and stromal features, including a collagen type III–rich ECM niche that sustains dormancy in the lung and restores proliferation when its architecture and density change [36], and astrocyte-deposited laminin-211 that promotes dormancy in the brain [37]. Dormancy is further shaped by cellular components of metastatic niches, including immune regulation in the liver [38], lung-resident alveolar macrophages that regulate the timing of breast cancer metastasis in the lung [39], and bone marrow mesenchymal stem cell niches that promote disseminated breast cancer cell dormancy via TGF-*β*2 and BMP7 signalling [40]. Together, these observations motivate the view that dormancy-to-outgrowth transitions depend on the coupled dynamics of cancer cell populations and their surrounding microenvironment.

Such coupling naturally suggests a collective perspective in which qualitative disease trajectories emerge from interactions between dormant cancer cells and the microenvironment, rather than from purely cell-intrinsic programs [41–43]. Feedback between populations and their shared microenvironment can generate threshold-like switches, multistability, and history dependence [44, 45]. Similar hysteretic memory logic appears in biological fate-control circuits, from developmental decisions to cancer cell-state plasticity [46, 47]. Cancer provides concrete examples of collective pathogenicity in which collective behaviour arises from cooperative or competitive interactions among cell populations and their microenvironment [48]. We therefore hypothesize that awakening from dormancy can emerge collectively from cancer–microenvironment coupling, and that a minimal representation of this coupling can already reproduce the qualitative dormancy behaviours observed experimentally and clinically.

Mathematical and computational modeling provides a quantitative way to formalize this hypothesis and to derive experimentally testable predictions [49, 50]. A broad spectrum of cancer modeling approaches exists [51, 52], ranging from mechanistic and multiscale simulations [53] to data-driven and treatment-optimization frameworks [54]. However, for dormancy and awakening the central challenge is not to reproduce every mechanistic detail, but to identify the minimal feedback logic that can generate distinct qualitative outcomes and to translate that logic into measurable parameters and perturbations. Evolutionary game theory (EGT) is well suited for this purpose because it captures frequency-dependent interactions and population-level adaptation in heterogeneous cellular communities, providing a compact framework to describe competition, coexistence, and takeover [55], while reducing complex microenvironmental influences to interpretable interaction parameters.

Despite extensive use of EGT in cancer [56–60], most existing models treat the microenvironment implicitly or as a fixed context, rather than as a dynamical state that is modified by cancer cells and that in turn could reshape cellular interactions. Several theoretical frameworks have begun to integrate ecological and environmental feedbacks into evolutionary dynamics [61, 62], yet explicit, experimentally interpretable couplings between cancer population dynamics and a primed microenvironment remain comparatively underdeveloped in the cancer dormancy setting. In addition, spatial structure and gradients can qualitatively alter evolutionary outcomes relative to well-mixed, mean-field dynamics [63–66]. While spatial effects are important in metastatic niches, a minimal well-mixed theory that isolates the core feedback mechanism provides a useful reduced model with clear, testable control parameters.

Here we present an EGT-based framework that couples interactions between cancer and host cells to an explicit conditioning factor as a microenvironmental state variable. Cancer cells produce this factor, which is cleared by the microenvironment and tolerated only up to a threshold, thereby reshaping the effective interaction landscape of the cancer–host system. This feedback makes dormancy dynamic and allows dormancy-to-outgrowth transitions to emerge from microenvironmental change. We show that this minimal coupling is sufficient to reproduce key qualitative dormancy behaviours as distinct longterm regimes. By linking the conditioning factor level to the interaction payoffs, microenvironmental priming can shift the effective competition between cancer and host cells, generating an environmental load threshold and, in the priming scenario, creating an initial-condition boundary that produces history-dependence in the transition from dormancy to outgrowth. Moreover, the framework reduces the system to a small set of experimentally interpretable parameters: (i) the environmental load combines the production, clearance, and tolerance of the conditioning factor into a single quantity that determines whether the factor can exceed the tolerance scale and strongly shift the microenvironment state, and (ii) interaction parameters summarize how the microenvironmental state biases cancer–host competition. This links qualitative dormancy behaviours to experimentally controllable processes and measurable readouts, enabling quantitative tests of the predicted transitions and guiding experiments on how conditioning factor production, clearance, or tolerance influence awakening.

## 2 Results

### 2.1 Replicator dynamics

We consider a well-mixed population composed of *S* strategies (cell types). Let *N*_*i*_ denote the abundance of strategy *i* and 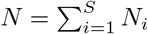 the total population size. The corresponding frequency is

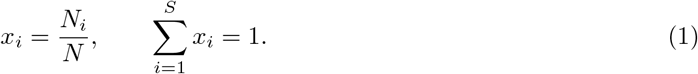

Interactions between cell types are encoded by a payoff matrix *M* ∈ ℝ^*S*×*S*^, where *M*_*ij*_ is the payoff (per-capita growth contribution) to a cell of type *i* interacting with a cell of type *j*. In a well-mixed population, the expected payoff (fitness) of strategy *i* is

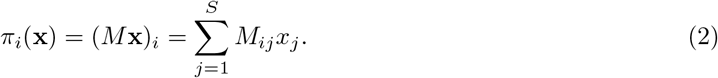

Assuming growth with rate *π*_*i*_(**x**),

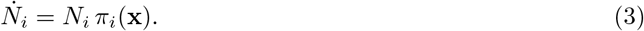

The mean population payoff is

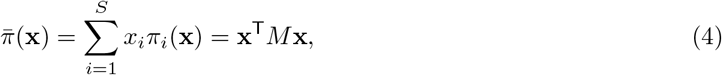

and differentiating *x*_*i*_ = *N*_*i*_*/N* (see Supplementary Section 1.1) we get the replicator equation

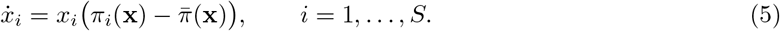

In our case, we assume there are two strategies or cell types, which are host cells (strategy 1), representing the non-cancer cellular component of the microenvironment, and cancer cells (strategy 2). Let *x*_2_ be the cancer fraction and *x*_1_ = 1 − *x*_2_ the host fraction. For the payoff matrix

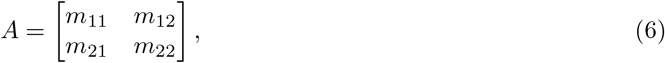

the replicator equation 5 for *x*_2_ becomes

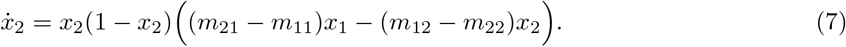

We define the parameters

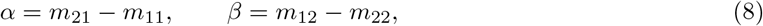

which quantify payoff differences. Specifically, *α* measures the advantage of being a cancer cell rather than a host cell when interacting with a host opponent (*j* = 1), while *β* measures the advantage of being host rather than cancer when interacting with a cancer opponent (*j* = 2). Equation 7 then reduces to

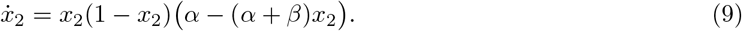

Using the invariance of replicator dynamics to baseline payoff shifts, it is convenient to work with the reduced matrix

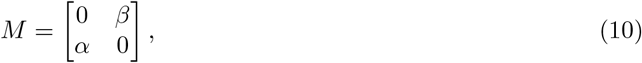

which is dynamically equivalent to *A* for the purposes of analyzing 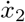 (see Supplementary Section 1.2). For readability, in what follows we denote the host fraction by *x*_*h*_ ≡ *x*_1_ and the cancer fraction by *x* ≡ *x*_2_. Thus, so *x*_*h*_ = 1 − *x*.

### 2.2 Baseline game without environmental feedback

We start from the simplest well-mixed, two-strategy model in which interaction incentives are constant in time. As a representative metastatic niche, and consistent with the schematic in Fig. 1A, we interpret the host compartment in the context of the bone marrow, a common site of breast cancer metastasis [12, 67]. This niche imposes distinct constraints on dormancy and outgrowth through differences in stromal composition, immune activity, and extracellular matrix properties [17]. The baseline replicator dynamics are

**Figure 1:**
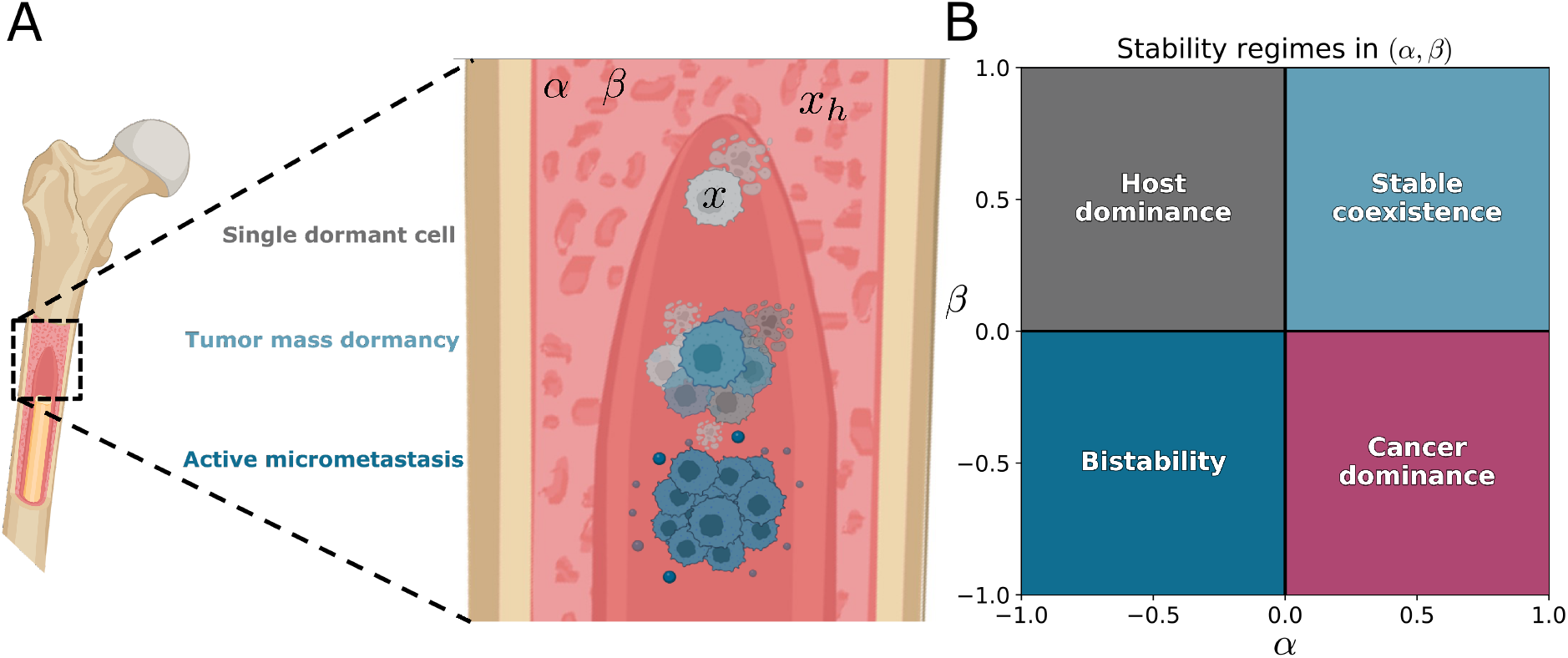
Baseline metastatic niche setting and static fate map for the two-strategy replicator model. **(A)** Schematic of disseminated cancer cells in a metastatic niche, illustrated as the bone marrow, highlighting clinically silent states across scales: single-cell dormancy, tumor-mass dormancy, and subclinical active micrometastasis. In the baseline model, the cancer fraction is *x* and the host fraction is *x*_*h*_ = 1 − *x*, interacting through constant payoff-difference parameters *α* and *β*. **(B)** Stability regimes in the (*α, β*) plane for Eq. (11). Host dominance, *α* < 0 and *β* > 0, converges to the boundary fixed point *x*\* = 0. Cancer dominance, *α* > 0 and *β* < 0, converges to the boundary fixed point *x*\* = 1. Stable coexistence, *α* > 0 and *β* > 0, converges to the interior fixed point *x*\* = *α/*(*α* + *β*). Bistability, *α* < 0 and *β* < 0, has two stable boundary fixed points at *x*\* = 0 and *x*\* = 1, separated by an unstable interior fixed point.

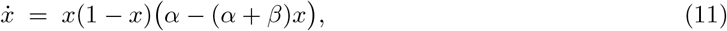

where *x* ∈ [0, 1] is the cancer fraction and (*α, β*) summarize payoff differences between cancer and host cells. We treat microenvironment-specific effects in a coarse-grained way through the interaction parameters (*α, β*). Since *α* and *β* are constant, Eq. (11) defines a single game for the whole experiment. Figure 1A shows this baseline model in the context of clinically silent metastatic disease. Here, single dormant DCC, tumor-mass dormancy, and active micrometastasis refer to different asymptomatic configurations across scales [17]. In the baseline model we do not represent these as distinct phenotypic states; instead, we use the cancer fraction *x* as an aggregate variable, and interpret the long-term fates of *x* as qualitative outcomes that can be mapped onto these configurations.

Equation (11) has three candidate fixed points,

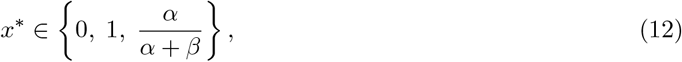

with an admissible interior fixed point only when *α* · *β* > 0 (see Supplementary Section 2). This produces four qualitatively distinct stability regimes in the (*α, β*) plane (Figure 1B): host dominance (*x* → 0) for *α* < 0, *β* > 0, cancer dominance (*x* → 1) for *α* > 0, *β* < 0, stable coexistence with a stable interior fixed point for *α* > 0, *β* > 0, and bistability for *α* < 0, *β* < 0, where both boundaries are stable and the interior fixed point is unstable, so trajectories go to *x* = 0 or *x* = 1 depending on the initial condition. This is illustrated in Fig. 2, where trajectories evolve in time but, for fixed (*α, β*), they always approach the same fixed points for a given choice of (*α, β*).

**Figure 2:**
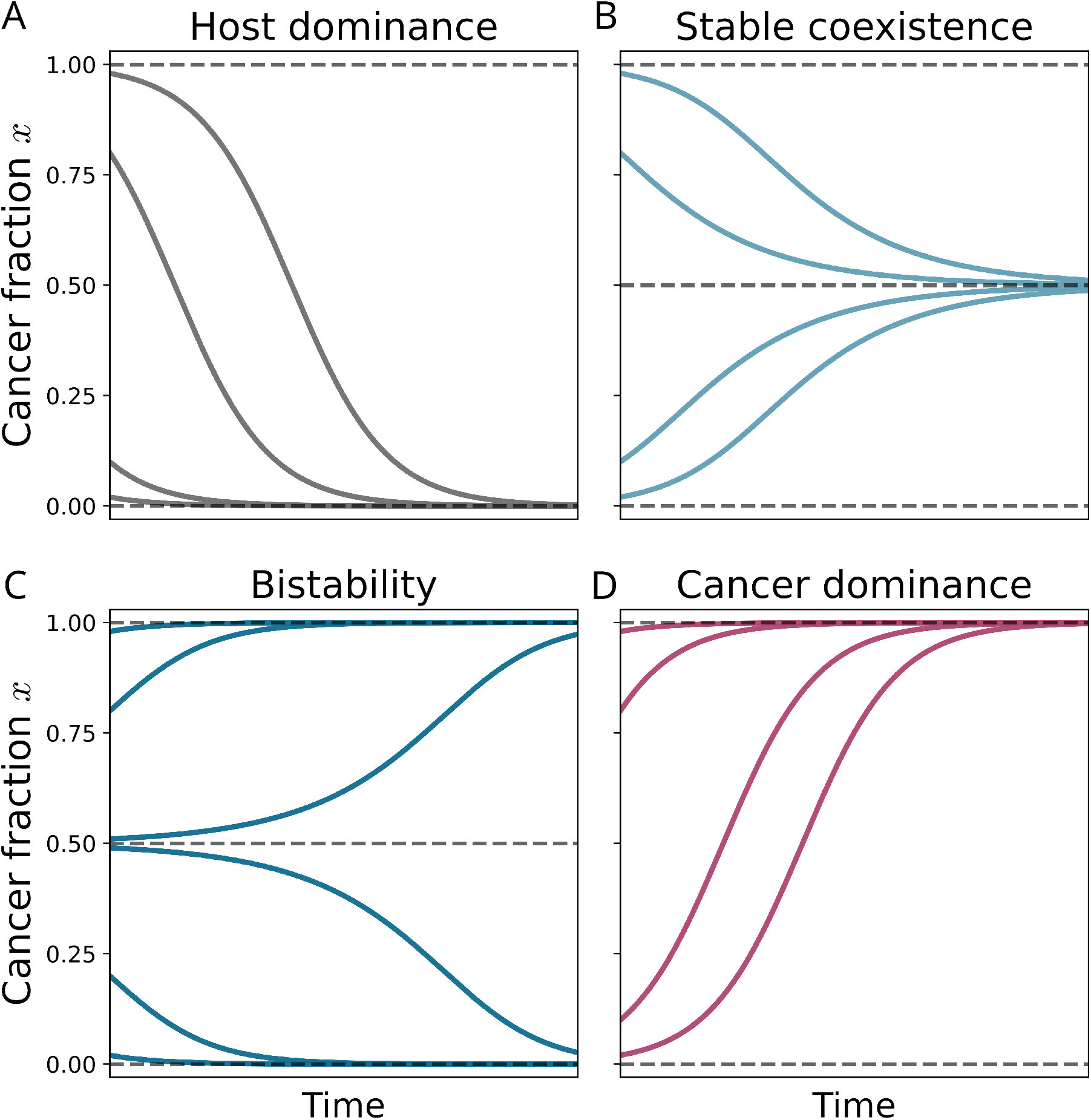
Baseline replicator dynamics converge to fixed points determined by constant. (*α, β*). Representative trajectories of the cancer fraction *x*(*t*) for Eq. (11). In each panel, (*α, β*) are fixed and only the initial condition *x*(0) varies. Parameters are: **(A)** Host dominance, (*α, β*) = (−0.5, +0.5), shown for *x*(0) ∈ {0.02, 0.10, 0.80, 0.98}, trajectories converge to the boundary fixed point *x*\* = 0. **(B)** Stable coexistence, (*α, β*) = (+0.5, +0.5), shown for *x*(0) ∈ {0.02, 0.10, 0.80, 0.98}, trajectories converge to the interior fixed point *x*\* = *α/*(*α* + *β*) = 0.5. **(C)** Bistability, (*α, β*) = (−0.5, *−*0.5), shown for *x*(0) ∈ {0.02, 0.20, 0.49, 0.51, 0.80, 0.98}, trajectories converge to *x*\* = 0 or *x*\* = 1 depending on whether *x*(0) is below or above the unstable interior fixed point *x*\* = *α/*(*α* + *β*) = 0.5. **(D)** Cancer dominance, (*α, β*) = (+0.5, −0.5), shown for *x*(0) ∈ {0.02, 0.10, 0.80, 0.98}, trajectories converge to the boundary fixed point *x*\* = 1. Dashed lines mark *x*\* = 0, *x*\* = 1, and, when present, the interior fixed point.

This baseline provides a compact and interpretable classification of long-term fates, but it is intrinsically static. For fixed (*α, β*), the attractor structure is fixed, and the system retains the same fixed points and basins of attraction, that is, the sets of initial conditions that converge to them, throughout the experiment (Fig. 2). As a result, obtaining a different qualitative outcome requires changing (*α, β*) externally rather than having transitions emerge from the system dynamics. As a consequence, the baseline model cannot account for fate switching driven by progressive microenvironmental changes, nor does it provide a mechanistic way to relate delayed transitions to measurable microenvironmental processes. This limitation motivates introducing state-dependent interactions via an explicit conditioning factor as a state variable, so that the effective game evolves with the microenvironment and qualitative transitions can arise endogenously.

### 2.3 Game with environmental feedback

Here we ask whether a minimal microenvironmental coupling can reproduce qualitative behaviours associated with disseminated cancer cell (DCC) dormancy and outgrowth. We define dormancy as a stable coexistence state, awakening as the loss of that stability, and outgrowth as the subsequent increase in cancer burden. In the scenarios considered here, awakening leads to convergence to cancer dominance as microenvironmental conditions cross a control threshold. To capture this mechanism, we next introduce an explicit coupling between cell competition and microenvironmental priming. Here, we use priming to refer to the history-dependent modification of the microenvironment by prior cancer cell presence, mediated through accumulation of the conditioning factor. In the model, this corresponds to an increase in the conditioning factor level 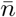. Biologically, we consider DCCs arriving to a niche such as the bone marrow (Figure 3A). We represent the cancer fraction by *x* ∈ [0, 1] and the host fraction by *x*_*h*_ = 1 − *x*. As in the baseline model, frequency-dependent competition is summarized by two payoff-difference parameters: *α* quantifies the advantage of cancer over host in interactions with host cells, while *β* quantifies the advantage of host over cancer in interactions with cancer cells.

**Figure 3:**
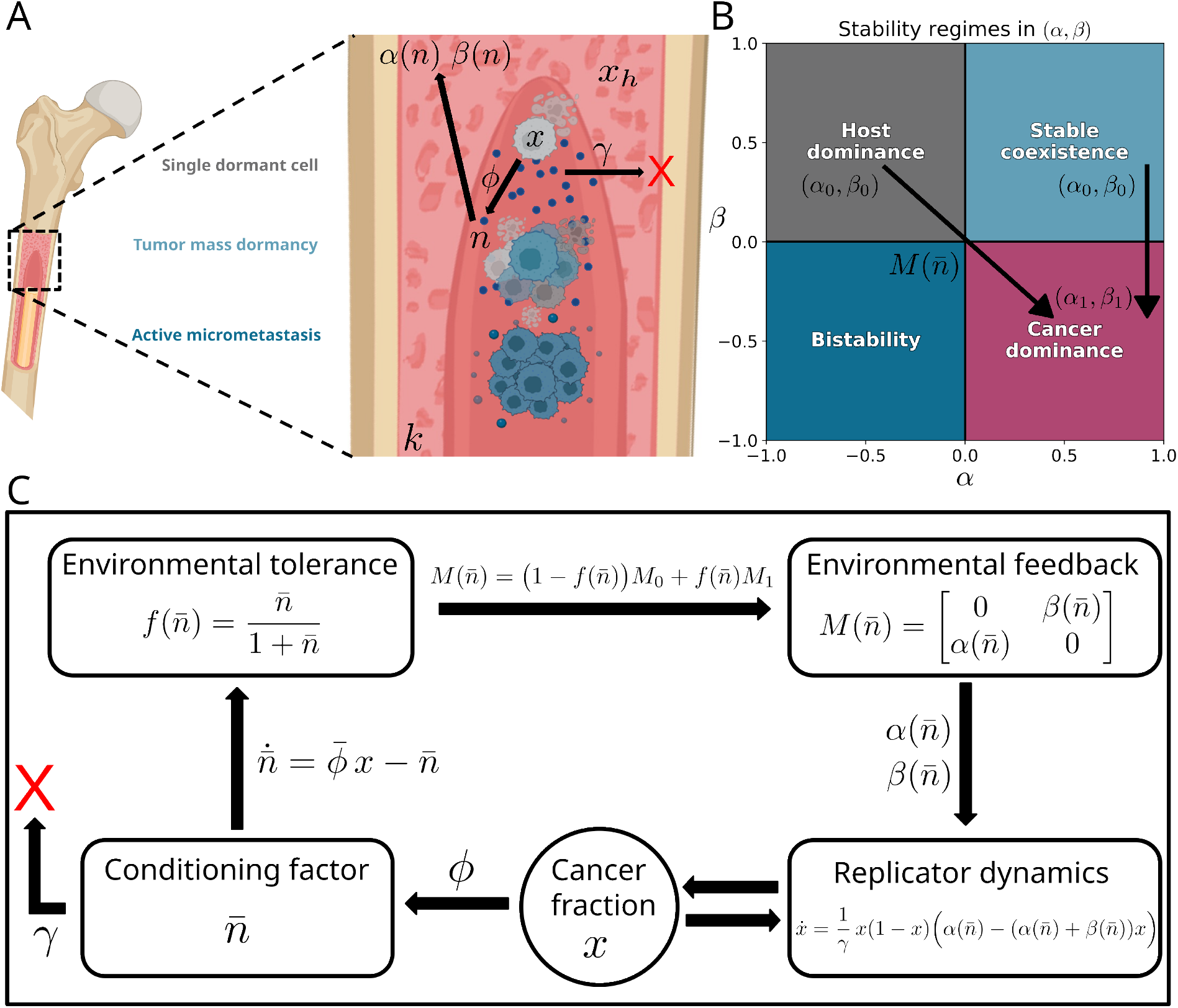
Feedback couples cell competition to microenvironmental priming. **(A)** Schematic of disseminated cancer cells in a metastatic niche, illustrated in the bone marrow. **(B)** Stability regimes in the (*α, β*) plane, as in Fig. 1B. In the feedback model, the conditioning factor level 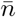 changes the effective interaction parameters, so the effective game moves through this plane as 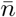 evolves. The arrows indicate two representative endpoint choices used later in this work: a colonization and priming scenario, in which the effective game shifts from the host-dominant quadrant (*α*_0_, *β*_0_) to the cancerdominant quadrant (*α*_1_, *β*_1_), and a dormancy-to-outgrowth scenario, in which the effective game shifts from the stable coexistence quadrant (*α*_0_, *β*_0_) to the cancer-dominant quadrant (*α*_1_, *β*_1_). **(C)** Dynamic dormancy theoretical framework based on EGT: the cancer fraction *x* competes with the host fraction *x*_*h*_ = 1 − *x* via frequency-dependent interactions. Cancer cells produce a conditioning factor *n* at rate *ϕ*, while the microenvironment clears it at rate *γ* and tolerates it up to a characteristic scale *k*. Using 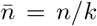, a tolerance gate 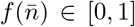 interpolates between two endpoint games, 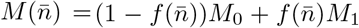. This defines the effective parameters 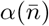 and 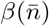 and closes the feedback loop by updating the competition dynamics of *x*.

To model microenvironmental priming, we introduce a scalar conditioning factor *n* ≥ 0 that is produced by cancer cells and cleared by the microenvironment (e.g. a generic signal standing in for cytokines, proteases, or remodeling enzymes). Cancer cells produce the conditioning factor *n* with production rate parameter, yielding a total production term proportional to the cancer fraction *x*, while the microenvironment clears it with rate *γ*, proportional to the current amount of conditioning factor. The microenvironment can tolerate the conditioning factor only up to a characteristic scale *k*, which sets the threshold between low- and high-priming microenvironmental conditions (Figures 3A and 3C). We work with the nondimensional conditioning factor level 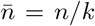 and measure time in clearance units *τ* = *γt* (see Supplementary Section 3).

Feedback enters by allowing the interaction incentives *α* and *β* to depend on the conditioning factor level 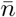, that is, the effective competition parameters become 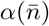 and 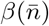 (Figure 3B). As 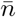 increases or decreases, the system changes the effective game it is playing while the population fractions evolve. In contrast to approaches that treat the environment as a bounded abstract variable and interpolate payoffs directly [61, 62], we model the microenvironment as a conditioning factor produced by cancer cells and cleared by the microenvironment, and we couple it back to the game through a nonlinear tolerance gate 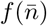.

Environmental priming modulates the effective interaction game by interpolating between two endpoint regimes. We define *M*_0_ as the low-priming (baseline) microenvironment game, corresponding to small conditioning factor levels, and *M*_1_ as the high-priming (primed) microenvironment game, corresponding to elevated conditioning factor levels. For two strategies we write these endpoint games in reduced form as

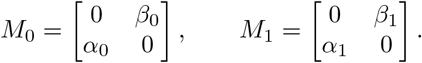

As the conditioning factor level increases, the effective game moves through the (*α, β*) plane along a trajectory that connects (*α*_0_, *β*_0_) to (*α*_1_, *β*_1_) (Fig. 3B). The arrows in Fig. 3B are schematic paths in the (*α, β*) plane, and they represent the two endpoint choices analyzed in the next section. We model this by the interpolation

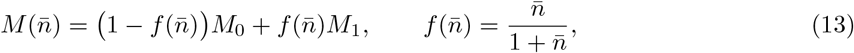

and the corresponding effective payoff-difference parameters are

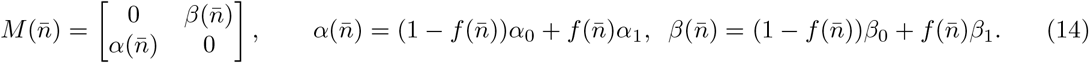

We discuss representative choices of *M*_0_ and *M*_1_ as microenvironment hypotheses in the next section.

Thus, the coupled system is

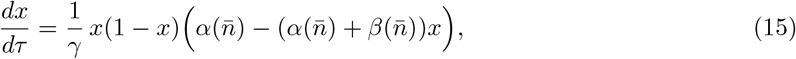

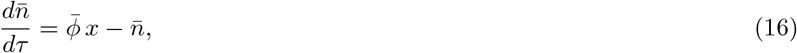

where 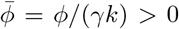 is the dimensionless effective production rate (environmental load). At *x* = 1, the steady state of Eq. (16) is 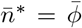, so 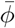 controls whether trajectories can reach the high-priming regime and, more generally, the speed and extent of microenvironmental priming (see Supplementary Section 3 for further details). Equivalently, the conditioning factor shifts the effective game from (*α*_0_, *β*_0_) toward (*α*_1_, *β*_1_) as 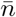 increases, while the population state *x* determines how rapidly the microenvironment becomes primed.

This coupling adds three main features. First, the dynamics can move through the (*α, β*) plane as the conditioning factor accumulates or clears, rather than remaining fixed in a single baseline regime (Figure 3B). Second, interactions become state-dependent, as 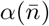 and 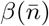 change because 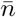 changes, and 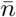 changes because *x* changes (Figure 3C). Third, this creates a mechanistic route from measurable microenvironmental processes, production rate *ϕ*, clearance rate *γ*, and tolerance scale *k*, to qualitative fate changes, enabling dormancy-to-awakening transitions to emerge from the system dynamics instead of requiring externally chosen (*α, β*).

### 2.4 Minimal coupling recapitulates dynamic dormancy behavior

Using the feedback model above, we now characterize the qualitative outcomes that emerge as the conditioning factor level changes and the microenvironment becomes primed. The coupled dynamics are controlled by the environmental load 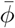, which determines whether conditioning factor remains buffered 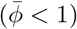 or can exceed the tolerance scale 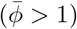.

Here we use the endpoint games *M*_0_ and *M*_1_ defined above, corresponding to low and high priming, respectively, to characterize how priming changes the effective interaction parameters (Eq. (14); Supplementary Section 4). To illustrate how endpoint choices shape qualitative behaviour, we analyze two representative microenvironment hypotheses (Supplementary Section 4), shown schematically by the two arrows in Fig. 3B. These arrows represent paths of the effective interaction parameters in the (*α, β*) plane as the conditioning factor level 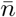 changes. The first is a colonization and priming scenario in which *M*_0_ lies in the host-dominant quadrant, *α*_0_ *<* 0 and *β*_0_ *>* 0, while *M*_1_ lies in the cancer-dominant quadrant, *α*_1_ *>* 0 and *β*_1_ *<* 0. The second is a dormancy-to-outgrowth scenario in which *M*_0_ lies in the coexistence quadrant, *α*_0_ *>* 0 and *β*_0_ *>* 0, while *M*_1_ again lies in the cancer-dominant quadrant, *α*_1_ *>* 0 and *β*_1_ *<* 0. These two endpoint choices underlie the four qualitative outcomes shown in Fig. 4. The host-dominant to cancer-dominant path underlies clearance and priming and multi-wave seeding, whereas the coexistence to cancer-dominant path underlies stable dormancy and awakening.

**Figure 4:**
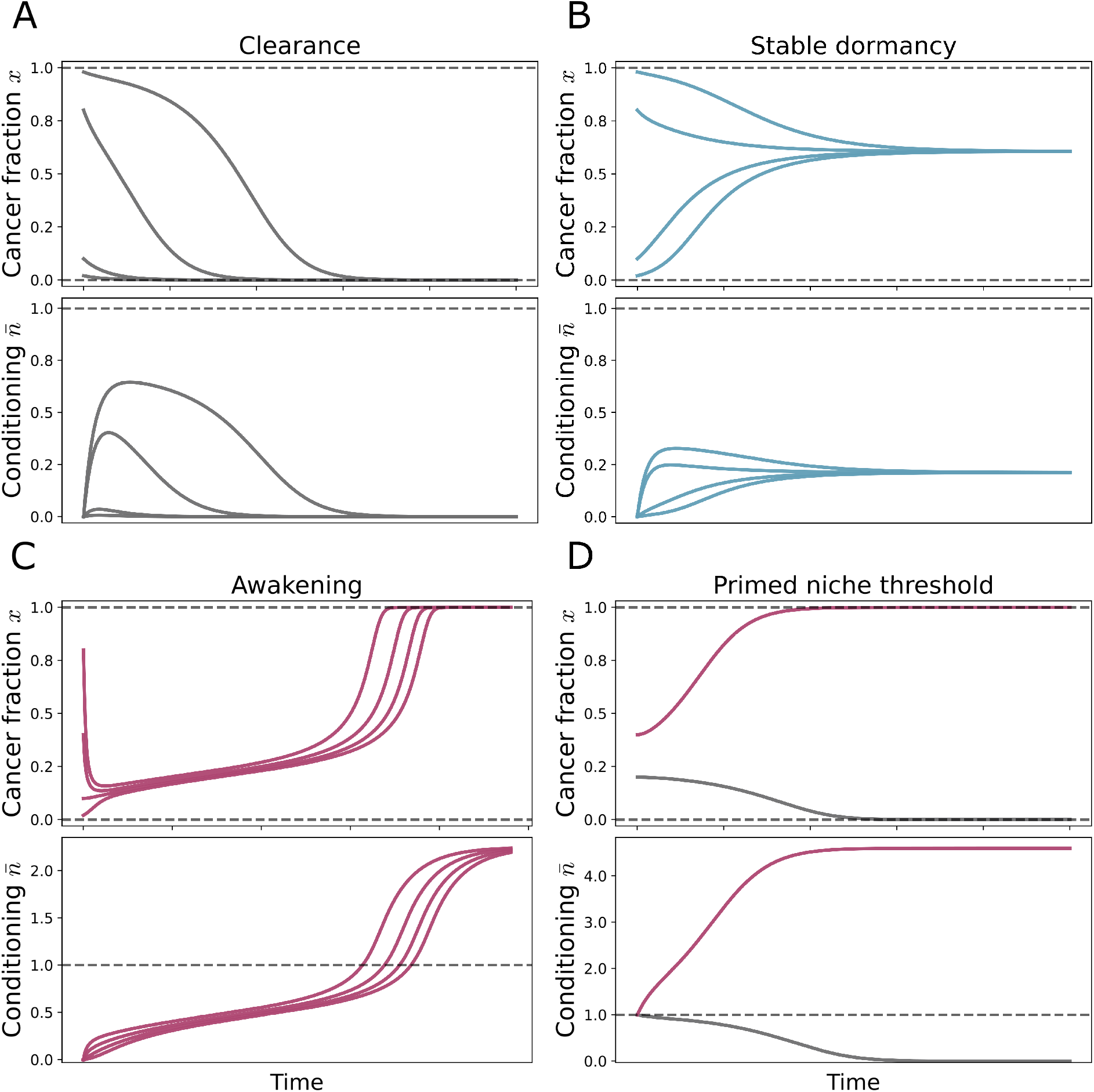
Minimal feedback generates clearance, stable dormancy, awakening, and priming. Simulations of Eqs. (15)–(16) in clearance-time units *τ* = *γt*, showing the cancer fraction *x*(*τ*) (top row in each panel) and conditioning factor level 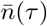 (bottom row in each panel). Dashed lines mark the boundary fixed points *x* = 0 and *x* = 1 and the tolerance scale 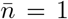. In each panel, the endpoint games are fixed by (*α*_0_, *β*_0_) and (*α*_1_, *β*_1_) and only initial conditions vary as indicated. **(A)** Clearance, host-dominant *M*_0_ to cancer-dominant *M*_1_ with (*α, β*) = (0.25, 0.5), *ϕ* = 0.7, *γ* = 0.5, *k* = 2.0, shown for *x*(0) ∈ {0.02, 0.10, 0.80, 0.98} and 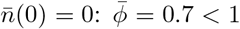 so the conditioning factor remains buffered below tolerance and *x*(*τ*) → 0 after a transient 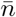 pulse. **(B)** Stable dormancy, coexistence *M*_0_ to cancerdominant *M*_1_ with (*α, β*) = (0.5, 0.5), *ϕ* = 0.7, *γ* = 1.0, *k* = 2.0, shown for *x*(0) 0.02, 0.10, 0.80, 0.98 and 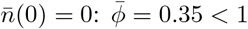 and trajectories converge to a stable interior fixed point with 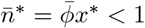. (**C**) Awakening, coexistence *M*_0_ to cancer-dominant *M*_1_ with (*α, β*) = (0.1, 0.9), *ϕ* = 0.09, *γ* = 0.02, *k* = 2.0, shown for *x*(0) ∈ {0.02, 0.10, 0.40, 0.80} and 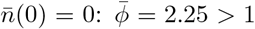 so the conditioning factor level can exceed tolerance and trajectories transition toward cancer dominance with *x*(*τ*) → 1. **(D)** Priming and multi-wave seeding, host-dominant *M*_0_ to cancer-dominant *M*_1_ with (*α, β*) = (0.5, 0.6), *ϕ* = 0.23, *γ* = 0.5, *k* = 0.1, shown for *x*(0) ∈ {0.20, 0.40} and 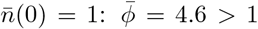 and outcomes depend on the initial state 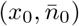, creating a threshold in initial conditions separating clearance from cancer dominance.

Unlike the baseline model with fixed (*α, β*), introducing feedback makes the effective game statedependent through 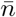, so the system can transition between interaction regimes without external parameter changes. This enables thresholds in initial conditions and history-dependence that cannot occur in the one-dimensional baseline dynamics. Figure 4 summarizes **four qualitative outcomes** of Eqs. (15)–(16) as representative time courses, and Table 1 provides the corresponding qualitative parameter requirements. The model equations are unchanged across panels; only parameter values and, when relevant, initial conditions differ. In the next section, we use phase portraits in the 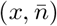 plane to explain the underlying fixed points and how their arrangement partitions the state space into different outcomes. The four outcome classes summarized in Fig. 4 and Table 1 are interpreted below in biological and dynamical terms.

**Table 1:** Summary of qualitative outcomes in the feedback model. Here 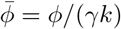 is the dimensionless environmental load, where *ϕ* is the production rate of the conditioning factor, *γ* is the clearance rate of the conditioning factor by the microenvironment, and *k* is the tolerance scale of the conditioning factor.

| Outcome | Environmental load $\bar{\phi}$ | Qualitative requirement on low- vs high-priming interactions |
| --- | --- | --- |
| Clearance | $< 1$ | Low-priming endpoint $M_0$ is host-favouring. The conditioning factor level remains buffered below tolerance, and trajectories converge to $x^* = 0$. |
| Stable dormancy | $< 1$ | Low-priming endpoint $M_0$ supports stable coexistence. The interior fixed point satisfies $\bar{n}^* = \bar{\phi}x^* < 1$, so coexistence is maintained. |
| Awakening | $> 1$ | The conditioning factor level can exceed tolerance and the effective interaction shifts toward the high-priming endpoint $M_1$ where cancer dominance is attracting. |
| Priming and multi-wave seeding | $> 1$ | The conditioning factor can persist. In the colonization and priming endpoint choice, outcomes can depend on the initial state $(x_0, \bar{n}_0)$, where $\bar{n}_0$ reflects prior microenvironmental priming, separating clearance from cancer dominance. |

#### Clearance

A small cancer fraction arrives and transiently increases the conditioning factor level, but the conditioning factor remains buffered below tolerance and *x*(*τ*) → 0 (Fig. 4A). In this outcome, 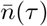 shows a short transient pulse and then returns toward 0. Clearance occurs for 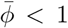 together with a low-priming interaction regime *M*_0_ that is host-favouring.

#### Stable dormancy

A small cancer fraction increases initially and then settles to a nonzero coexistence level without macroscopic outgrowth (Fig. 4B). In the model this corresponds to convergence to a stable interior fixed point *x*\* ∈ (0, 1), with 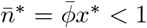 when 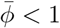. This behaviour requires that the low-priming endpoint *M*_0_ supports stable coexistence rather than clearance or cancer dominance.

#### Awakening

When 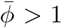, the conditioning factor level can exceed the tolerance scale and *x*(*τ*) can show a delayed increase followed by a transition to cancer dominance (Fig. 4C). Here, trajectories begin near a dormancy-like composition, but as 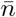 increases the effective interaction shifts toward the high-priming regime and the dynamics cross into cancer dominance, with *x*(*τ*) → 1.

#### Priming and multi-wave seeding

For 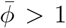, the same parameter values can lead to different outcomes depending on the initial cancer burden and whether the microenvironment is already primed (Fig. 4D). In this case, the coupled 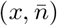 dynamics create a threshold in initial conditions: trajectories starting below it are cleared, whereas trajectories starting above it progress to cancer dominance. Biologically, this captures priming across waves, where earlier DCC arrivals elevate 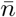 and shift the effective interaction so that later arrivals succeed under conditions that would otherwise lead to clearance.

In this model, two distinct thresholds appear. The environmental load 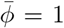 determines whether the conditioning factor can exceed the tolerance scale 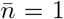 (that is, *n* = *k*) at full cancer burden, *x* = 1.

In addition, for the colonization and priming endpoint choice, in which *M*_0_ host-dominant and is *M*_1_ cancer-dominant, and for 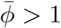, the dynamics introduce a threshold in state: for fixed parameters, the outcome can depend on whether the initial state 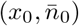 lies on one side or the other of a boundary in the 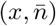 plane. Biologically, this means that two microenvironments with the same current cancer burden *x*_0_ can have different fates if their conditioning factor histories differ, reflected in different initial levels 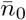. In this sense, priming denotes microenvironmental history dependence, where prior cancer cell presence elevates the conditioning factor level 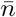 and shifts the effective interaction parameters toward the high-priming endpoint, enabling later DCC arrivals to succeed under conditions that would otherwise lead to clearance in an unprimed microenvironment [9].

Across all four outcomes, feedback makes interactions state-dependent: 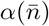 and 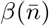 change because 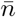changes, and 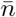 changes because *x* changes. This closes a loop in which the cancer cell population *x* modifies the microenvironment and the modified microenvironment reshapes competition, so transitions such as dormancy-to-outgrowth can arise endogenously without externally changing (*α, β*). In the next section, we use phase portraits in the 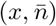 plane to show how this coupling reshapes the fixed points, that is, the long-term states of the system, and the basins of attraction, that is, the regions of initial conditions that converge to them.

### 2.5 Phase portraits show how feedback changes outcomes endogenously

Time traces in Fig. 4 show the four qualitative outcomes of the feedback model: clearance, stable dormancy, awakening, and priming and multi-wave seeding. To explain why these outcomes occur, we visualize the coupled dynamics in the 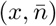 plane (Fig. 5). The key point is that the conditioning factor level 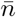 is a dynamical variable, not a fixed parameter, as it increases with cancer burden and decreases through microenvironmental clearance. Because 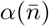 and 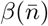 depend on 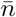, the effective interaction changes along trajectories, so transitions can occur without externally changing parameters. In regimes with more than one stable long-term state (Fig. 5D), the phase portrait also reveals a boundary in initial conditions, which is the source of history-dependence and underlies microenvironmental priming.

**Figure 5:**
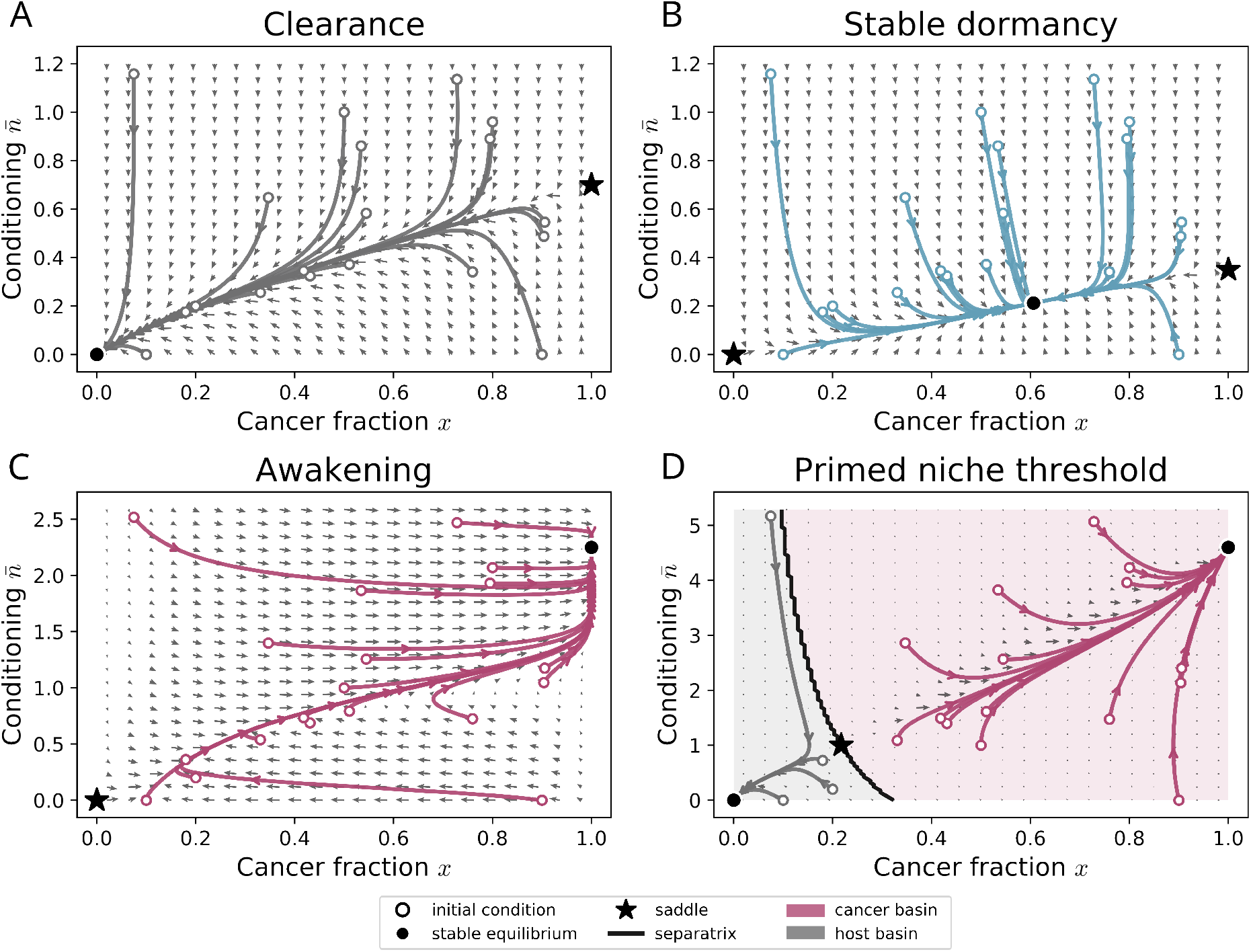
Phase portraits show how microenvironmental feedback reshapes long-term outcomes. Vector fields and representative trajectories of Eqs. (15)–(16) in the 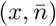 plane. Open circles denote initial conditions and colored curves show the corresponding trajectories. Filled circles mark stable fixed points and stars mark saddle points. In panel D, the thick black curve denotes the basin boundary (separatrix), and shaded regions indicate the corresponding basins of attraction. In each panel, trajectories were computed from 19 representative initial conditions, including 14 randomly sampled initial conditions and 5 fixed reference initial conditions. **(A)** Clearance: host-dominant *M*_0_ to cancer-dominant *M*_1_ with (*α, β*) = (0.25, 0.5), *ϕ* = 0.7, *γ* = 0.5, *k* = 2.0, and 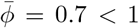. Trajectories converge to 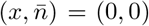. **(B)** Stable dormancy: coexistence *M*_0_ to cancer-dominant *M*_1_ with (*α, β*) = (0.5, 0.5), *ϕ* = 0.7, *γ* = 1.0, *k* = 2.0, and 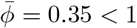. Trajectories converge to a stable interior fixed point 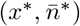. **(C)** Awakening: coexistence *M*_0_ to cancer-dominant *M*_1_ with (*α, β*) = (0.1, 0.9), *ϕ* = 0.09, *γ* = 0.02, *k* = 2.0, and 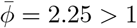. Trajectories converge to cancer dominance 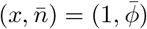. (**D**) Priming and multi-wave seeding: host-dominant *M*_0_ to cancer-dominant *M*_1_ with (*α, β*) = (0.5, 0.6), *ϕ* = 0.23, *γ* = 0.5, *k* = 0.1, and 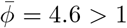. A saddle fixed point generates a basin boundary in initial conditions that separates clearance from cancer dominance, capturing priming as a history-dependent effect through the state variable 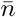.

In each panel of Fig. 5, arrows show the local direction of change of cancer fraction *x* and conditioning factor level 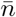, and colored curves show representative trajectories from different initial conditions (open circles). Filled circles mark stable fixed points, that is, long-term states approached by nearby trajectories, and stars mark saddle points, which attract trajectories in some directions and repel them in others. In panel D, the thick black curve denotes the basin boundary (separatrix) that partitions the 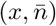 plane into regions leading to different long-term outcomes, and shaded regions indicate the corresponding basins of attraction. We now interpret each panel of Fig. 5 in terms of the long-term outcomes in Fig. 4 and the corresponding fixed points and basins in the 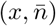 plane. The fixed points and their local stability for the two representative endpoint choices are summarized in the Supplementary Tables S1 and S2.

#### Clearance and stable dormancy correspond to a single dominant attractor for 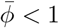

When 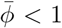, the conditioning factor level remains buffered below tolerance even at full cancer burden (*x* = 1), so trajectories stay in the low-priming range of 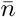. In the clearance panel (Fig. 5A), trajectories from small cancer fractions move toward the host-dominant state 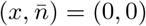, so cancer declines and the conditioning factor level returns toward 0 after a transient pulse. In the stable dormancy panel (Fig. 5B), the same buffering of 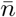 coexists with an interaction structure that supports an interior coexistence fixed point. As a result, trajectories converge to a stable interior fixed point 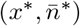, capturing dormancy without outgrowth. In other words, when *γk* is large relative to *ϕ*, the environmental load remains below threshold 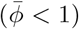, so the conditioning factor stays buffered and the system can support clearance or stable dormancy.

#### Awakening occurs for 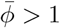 when priming can access the high-priming interaction regime

When 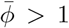, the conditioning factor level can exceed the tolerance scale 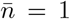 at sufficiently high cancer burden, so trajectories can enter the high-priming range of 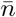 (see Supplementary Table S2 for the corresponding fixed-point structure and stability conditions). In the awakening panel (Fig. 5C), trajectories pass through a region where the effective interaction shifts toward cancer dominance at elevated 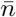. For some initial conditions, this transition is not monotonic, and the cancer fraction can decrease transiently before later increasing toward cancer dominance, reflecting a temporary dormancy-like or regression-like phase before awakening. As a result, trajectories converge to the cancer-dominant equilibrium 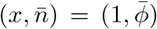, often with coupled changes in 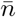 and *x*, as elevated 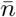 shifts the effective interaction toward cancer dominance.

#### Priming and multi-wave seeding arise from an initial-condition boundary in the priming scenario

A key qualitative addition of the coupled dynamics is the appearance of a basin boundary (separatrix) in initial conditions (Fig. 5D, black curve), generated by a saddle fixed point (see Supplementary Table S1). This boundary partitions the 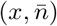 plane into two regions: initial states on one side converge to clearance 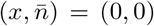, whereas initial states on the other side converge to cancer dominance 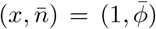. Biologically, this means that a microenvironment can clear an early DCC arrival when both the cancer fraction and conditioning factor level are low, but becomes more cancer permissive after prior priming, reflected in an elevated 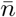. In that case, a later arrival can start on the cancer-dominant side of the boundary and progress to outgrowth without any external parameter change. In other words, priming is not imposed by switching the game externally; it emerges because the coupled system can encode microenvironmental history in the state variable 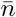.

## 3 Discussion

Dormant disseminated cancer cells can persist for long periods in distant microenvironments, yet the mechanisms that trigger their transition to metastatic outgrowth remain unclear. A central challenge is that dormancy and outgrowth are not simply different endpoints, but dynamic processes shaped by a changing microenvironment [68, 69]. This makes it difficult to connect qualitative clinical observations, including stable dormancy, delayed outgrowth, and microenvironmental priming, to quantitative variables that experiments can perturb and measure. Here we developed a theoretical framework that couples interactions between cancer and host cells to an explicit conditioning factor, *n*, as a microenvironmental state variable. In the baseline replicator model with fixed interaction parameters (*α, β*), the interaction game is fixed once (*α, β*) are specified, and so is the long-term outcome. This baseline provides a useful classification of outcomes, but it cannot generate endogenous switching from dormancy to outgrowth because there is no evolving microenvironmental state. Introducing feedback closes the loop between the cancer cell population state and microenvironmental state. The cancer fraction *x* drives the conditioning factor level 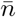, and 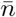 updates the effective interaction parameters 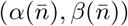. This minimal coupling is sufficient to generate endogenous state transitions without externally changing (*α, β*), including dormancy to outgrowth transitions that emerge from the dynamics of microenvironmental priming.

Two distinct thresholds organize the behaviour. First, the environmental load threshold 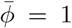 determines whether the conditioning factor can exceed the tolerance scale 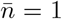 at full cancer burden, *x* = 1. This sets whether strong priming is reachable at all. Second, in the colonization and priming scenario, where *M*_0_ is host-dominant and *M*_1_ is cancer-dominant, the coupled dynamics generate a boundary in initial conditions when 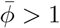. In that case, the outcome depends on where the initial state 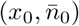 lies in the 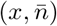 plane. This is the basis of priming and multi-wave seeding: two systems with the same initial cancer burden *x*_0_ can still have different fates if they differ in their initial conditioning factor level 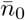, which reflects different microenvironmental histories and priming.

This interpretation is also consistent with previous game-theoretic work framing dormancy as a criticality problem in interacting host–cancer populations, where outcomes become highly sensitive to perturbations near unstable equilibria [70–73]. More broadly, the resulting tipping point dynamics is consistent with experimental observations in which small changes in initial burden or in the parameters controlling microenvironmental feedback lead to qualitatively different outcomes [74]. Relatedly, introducing novel interactions into the tumor microenvironment through secreted factors can non-cell-autonomously shift growth trajectories, supporting the view that interaction structure and feedback can generate counterintuitive population-level outcomes [75]. These ideas align with broader arguments that collective cell behaviour contributes to cancer progression and resistance, and that we still lack quantitative models that integrate the diversity of short-lived and long-lived interactions across scales [42, 43].

The same feedback logic is also relevant to premetastatic niche biology. Premetastatic niche studies emphasize systemic tumor-derived signals that remodel distant niches before overt colonization, thereby altering the probability of later establishment and growth [76]. Experimental work showing that breast cancer-secreted factors can systemically induce a fibrotic lung premetastatic niche provides a concrete instance of the same coupling, in which tumor burden increases a conditioning factor, and the modified microenvironment becomes more permissive for subsequent metastasis [77]. In our framework, such premetastatic priming can be represented either as a sustained source term for *n* or, more simply, as a primed initial condition 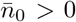 before DCC arrival, which is exactly the setting in which an initialcondition boundary becomes relevant. Recent work on nascent tumor persistence points in the same direction, showing that early lesions can diverge in fate depending on whether they induce a supportive stromal microenvironment [78]. These systems are clearly more complex than a two-variable model, but they reinforce the same question: which microenvironmental processes control conditioning factor accumulation, and which determine the direction and magnitude of the microenvironment-dependent changes in 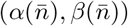?

A deliberate feature of the present model is that (*α, β*) are coarse-grained parameters. They summarize the net effect of multiple microenvironmental components on relative competitive advantage, including biochemical cues, immune activity, and biophysical constraints imposed by the ECM. This is useful because many perturbations act through several mechanisms simultaneously, and it motivates interpreting shifts in (*α, β*) as signatures of how the microenvironment biases competition. From a systems mechanobiology perspective, plausible contributors to these effective incentives include ECM composition and architecture, stiffness and viscoelasticity, ligand availability, adhesion-mediated mechanotransduction such as integrin–FAK signalling, and growth-limiting mechanical stress [79–83]. This interpretation is consistent with dormancy studies implicating cell–matrix interactions and organ-specific stromal or ECM states in maintaining or releasing dormant cells [35–38, 84]. Within our framework, such processes need not be modeled separately. Instead, they can be viewed as shifting the endpoint locations of *M*_0_ and *M*_1_ in the (*α, β*) plane and the speed and extent of priming through 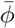. This suggests an experimental strategy in which one first estimates effective (*α, β*) under controlled microenvironmental perturbations, including ECM composition, stiffness, geometry, degradability, and stromal co-culture, and then interprets those shifts as coarse-grained readouts of how the microenvironment biases competition. Such controlled microenvironmental perturbations could also be implemented in biomaterial-based platforms that sustain dormant states, enabling tests of model predictions on dormancy versus awakening [85].

The model is intentionally minimal and therefore has clear limitations. It is well-mixed, deterministic, restricted to two cell types, and includes a single scalar microenvironmental variable, so it does not capture explicit spatial gradients, stochastic seeding and extinction, phenotypic switching among dormant and proliferative states, or heterogeneity in the microenvironmental response. Nevertheless, it remains valuable as a reduced theory that isolates a feedback mechanism and yields interpretable control parameters. One extension is to include additional strategies, for example separating dormant, proliferative, stromal, or immune-associated states, so that collective dormancy is represented as a heterogeneous population rather than a single cancer strategy [17, 86]. Another is to relax the well-mixed assumption. In most tissues, cells are spatially constrained and interact in a distance-dependent manner, and spatial structure can qualitatively alter evolutionary outcomes relative to mean-field dynamics [63–65], including under weak selection [87]. A further extension is to include stochasticity. Because metastatic seeding is sparse and early lesions are small, demographic fluctuations, random extinction, and rare escape events are likely to matter near regime boundaries. This is especially relevant because stochastic game structure remains underdeveloped in feedback formulations of evolutionary game theory [88], and spatial interactions can further modify strategy–environment feedback [66]. More realistic spatial extensions could also connect the theory more directly to experiments, for example, by resolving where stromal support arises and where resistant or awakened subpopulations emerge within tissue [89].

From a broader theoretical perspective, the model is a minimal instance of environment-mediated evolutionary dynamics in which strategies modify an environmental variable that then reshapes payoffs. This logic appears in ecological feedback games and eco-evolutionary models that couple replicator dynamics to a changing environment [61,62]. Our formulation differs in that the environmental variable is explicitly interpretable as a conditioning factor with production, clearance, and tolerance, rather than an abstract bounded environment state. This framing also connects naturally to niche construction theory, where organisms modify their selective environment and thereby change the subsequent evolutionary dynamics [90]. More broadly, environment-coupled game models have potential applications beyond dormancy, wherever populations reshape shared microenvironmental constraints such as cytokine fields, stromal activation, immune recruitment, or ECM remodeling, and where those changes feed back to alter competitive interactions [91, 92]. In this sense, the model is not only a description of dormancy, but also a general reduced framework for collective, feedback-driven state transitions in cancer and related multicellular systems.

In conclusion, our work identifies a minimal dynamical mechanism by which cancer cells modify the tumor microenvironment and the altered microenvironment feeds back on cancer-host competition, generating transitions between dormancy and outgrowth. This minimal coupling is sufficient to recapitulate multiple long-term regimes, including clearance, stable dormancy, awakening, and priming-dependent outgrowth, together with thresholded behaviour and history dependence. By reducing these dynamics to experimentally interpretable parameters linked to conditioning factor accumulation and microenvironment-dependent interaction changes, the framework provides a compact bridge between collective cancer dynamics and controllable stromal or extracellular matrix perturbations. More specifically, it provides a basis for asking which microenvironmental processes are sufficient to trigger dormant cancer cell awakening.

## 4 Methods

The baseline and feedback models analyzed in this study are introduced in Sections 2.2 and 2.3, with derivations and additional analytical details provided in the Supplementary Section 1.

### 4.1 Representative endpoint choices

To illustrate how feedback changes qualitative behaviour, we analyzed two representative endpoint choices. In the colonization and priming scenario, the low-priming endpoint *M*_0_ lies in the host-dominant quadrant and the high-priming endpoint *M*_1_ lies in the cancer-dominant quadrant. In the dormancy-to-outgrowth scenario, *M*_0_ lies in the coexistence quadrant and *M*_1_ again lies in the cancer-dominant quadrant. These endpoint choices were used as illustrative hypotheses for how microenvironmental priming reshapes cancer–host competition. Their analytical properties are detailed in Supplementary Section 4.

### 4.2 Phase plane analysis

Qualitative behaviour was analyzed from the fixed points of the baseline and feedback systems and from the geometry of trajectories in the 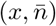 plane. For the feedback model, we identified stable and unstable equilibria and interpreted representative trajectories in terms of long-term regimes such as clearance, stable dormancy, awakening, and priming-dependent outgrowth. In the priming scenario, the phase portrait was also used to identify the boundary in initial conditions that separates clearance from cancer dominance. For phase portraits (Fig. 5), trajectories were initialized from 19 representative initial conditions, consisting of 14 randomly sampled initial conditions and 5 fixed reference initial conditions chosen to probe the boundary and interior regions of the phase plane. Analytical expressions for fixed points and their local stability are provided in the Supplementary Sections **??** and 4.

### 4.3 Numerical simulations and plotting

Numerical simulations were performed in Python using `NumPy` [93] for numerical computation, `SciPy` [94] for integration of the ordinary differential equations, and `Matplotlib` [95] for plotting. Time trajectories of the baseline and feedback models were obtained by numerically solving the corresponding systems of ordinary differential equations with `scipy.integrate.solve ivp`. Phase portraits were generated from the vector field of the feedback system together with representative trajectories from selected initial conditions.

Parameter values were chosen to illustrate the distinct qualitative regimes discussed in the text rather than to fit a specific experimental dataset. Figure-specific parameter values and initial conditions are reported in the corresponding figure captions. No parameter inference or calibration to experimental measurements was performed in this study.

### 4.4 Model variables and parameters

Table 2 summarizes the variables and parameters used in the model.

**Table 2:** Summary of variables and parameters used in the baseline and feedback models.

| Notation | Quantity | Description |
| --- | --- | --- |
| $x$ | Cancer fraction | Fraction of cancer cells in the population |
| $x_h = 1 - x$ | Host fraction | Fraction of host cells in the population |
| $n$ | Conditioning factor | Cancer-produced conditioning factor |
| $\bar{n} = n/k$ | Conditioning factor level | Conditioning factor scaled by tolerance |
| $\phi$ | Production rate | Rate of conditioning factor production by cancer cells |
| $\gamma$ | Clearance rate | Rate of conditioning factor removal by the microenvironment |
| $k$ | Tolerance scale | Characteristic tolerance scale of the conditioning factor |
| $\bar{\phi} = \phi/(\gamma k)$ | Environmental load | Composite measure of production, clearance, and tolerance |
| $\alpha$ | Payoff-difference parameter | Advantage of cancer over host against host |
| $\beta$ | Payoff-difference parameter | Advantage of host over cancer against cancer |
| $\alpha(\bar{n}), \beta(\bar{n})$ | Effective interaction parameters | Microenvironment-dependent competition parameters |
| $M_0$ | Low-priming endpoint game | Effective interaction regime at low conditioning factor level |
| $M_1$ | High-priming endpoint game | Effective interaction regime at high conditioning factor level |
| $f(\bar{n})$ | Tolerance gate | Interpolation between $M_0$ and $M_1$ |

## 5 Data availability

All the code used has been deposited in a publicly accessible Zenodo repository: https://doi.org/10.5281/zenodo.18925574 and in the GitHub repository https://github.com/gyanezfeliu/dynamics_dormancy.

## 6 Acknowledgments

G.Y.F. and A.C. would like to acknowledge funding from IKERBASQUE Basque Foundation for Science, from Fundación Científica Asociación Española Contra el Cáncer (grant LABAE223466CIPI), from the Spanish Ministry of Science and Innovation (MCIN/AEI/10.13039/501100011033/FEDER UE, through grant PID2021-123013OB-I00) and from the European Research Council Consolidator Grant (DORMATRIX, 101123883).

## 7 Author information

### 7.1 Authors and Affiliations

**Group of Bioengineering in Regeneration and Cancer, Biogipuzkoa Health Research Institute, San Sebastián, Spain**

Guillermo Yáñez Feliú, Amaia Cipitria.

**Department of Biomaterials, Max Planck Institute of Colloids and Interfaces, Potsdam, Germany**

Giacomo Rossato, Angelo Valleriani.

**IKERBASQUE, Basque Foundation for Science, Bilbao, Spain**

Amaia Cipitria.

### 7.2 Contributions

Conceptualization: G.R., A.V., A.C. Methodology: G.Y.F., G.R., A.V., A.C. Software: G.Y.F., G.R. Validation: G.Y.F., G.R., A.V., A.C. Formal analysis: G.Y.F., G.R. Investigation: G.Y.F., G.R. Resources: A.V., A.C. Data Curation: G.Y.F. Writing - Original Draft: G.Y.F. Writing - Review & Editing: G.Y.F., G.R., A.V., A.C. Visualization: G.Y.F. Supervision: A.V., A.C. Project administration: A.V., A.C. Funding acquisition: A.V., A.C. All authors read and approved the final manuscript.

